# Extensive DNA methylome rearrangement during early lamprey embryogenesis

**DOI:** 10.1101/2023.05.25.542242

**Authors:** Allegra Angeloni, Skye Fissette, Deniz Kaya, Jillian M. Hammond, Hasindu Gamaarachchi, Ira W. Deveson, Robert J. Klose, Weiming Li, Xiaotian Zhang, Ozren Bogdanovic

**Affiliations:** Garvan Institute of Medical Research, Sydney, Australia; School of Biotechnology and Biomolecular Sciences, University of New South Wales, Sydney, Australia; Department of Fisheries and Wildlife, Michigan State University, East Lansing, United States; Department of Biochemistry, University of Oxford, Oxford, United Kingdom; Genomics Pillar, Garvan Institute of Medical Research, Sydney, NSW, Australia; Centre for Population Genomics, Garvan Institute of Medical Research and Murdoch Children’s Research Institute, Australia; School of Computer Science and Engineering, University of New South Wales, Sydney, NSW, Australia; Faculty of Medicine, University of New South Wales, Sydney, NSW, Australia; Center for Epigenetics, Van Andel Research Institute, Grand Rapids, United States; University of Texas Health Science Center, Houston, TX, United States; Centro Andaluz de Biología del Desarrollo, CSIC-Universidad Pablo de Olavide-Junta de Andalucía Seville, Spain

**Keywords:** DNA methylation, epigenome evolution, embryogenesis, programmed genome rearrangement

## Abstract

DNA methylation (5-methylcytosine, 5mC) is a repressive gene regulatory mark widespread in vertebrate genomes, yet the developmental dynamics in which 5mC patterns are established vary across species. While mammals undergo two rounds of global 5mC erasure, the zebrafish genome exhibits localized maternal-to-paternal 5mC remodeling, in which the sperm epigenome is inherited in the early embryo. To date, it is unclear how evolutionarily conserved such 5mC remodeling strategies are, and what their biological function is. Here, we studied 5mC dynamics during the embryonic development of sea lamprey (*Petromyzon marinus*), a jawless vertebrate which occupies a critical phylogenetic position as the sister group of the jawed vertebrates. We employed base-resolution 5mC quantification in the lamprey germline, embryonic and somatic tissues, and discovered large-scale maternal-to-paternal epigenome remodeling that affects >30% of the embryonic genome and is predominantly associated with partially methylated domains (PMDs). We further demonstrate that sequences eliminated during programmed genome rearrangement (PGR), a hallmark of lamprey embryogenesis, are hypermethylated in sperm prior to the onset of PGR. Our study thus unveils important insights into the evolutionary origins of vertebrate 5mC reprogramming, and how this process might participate in diverse developmental strategies.

## INTRODUCTION

DNA methylation (5mC) is a chemical modification to the DNA that represents one of the most pervasive gene regulatory marks in vertebrates [1–3]. 5mC is predominantly found within the cytosine-guanine dinucleotide context (mCG), occurring at ∼80% of all CpG sites in vertebrate genomes [4–6]. mCG is associated with long-term silencing processes including somatic silencing of germline genes and silencing of repetitive DNA elements, as well as X-chromosome inactivation and genomic imprinting in mammals [7–10]. mCG is deposited by *de novo* DNA methyltransferases 3A/B (DNMT3A/B) and maintained following DNA replication by DNA methyltransferase 1 (DNMT1) [11–13]. DNA demethylation can occur passively via replication-coupled dilution [14,15], or actively via methylcytosine dioxygenase Ten-Eleven Translocation (TET) enzymes [16].

Mammalian development is characterized by two waves of global mCG erasure occurring in the preimplantation embryo and in the developing germline, followed by re-establishment of cell-type specific mCG [17–24]. However, it appears that such mCG remodeling processes are not evolutionarily conserved. For example, the zebrafish genome does not undergo global DNA demethylation [10,25–27]. Instead, the maternal methylome is remodeled to match the hypermethylated paternal genome prior to the onset of zygotic genome activation (ZGA) [28]. Thus, mCG patterns in the early embryo closely resemble those of sperm. Moreover, we have recently identified maternal-to-paternal DNA methylome remodeling in medaka embryos, revealing evolutionary conservation of developmental epigenome dynamics in distantly related teleost species [29].

Unlike hypermethylated vertebrate genomes, genomes of non-vertebrate lineages typically display “mosaic” mCG patterns, characterized by regions of heavily methylated DNA interspersed with fully unmethylated domains [3,30,31]. To date, it remains elusive how and why vertebrate genomes transitioned to hypermethylation as a default state. The sea lamprey *Petromyzon marinus* is an extant jawless fish that can serve as a valuable model organism for understanding mCG evolution in metazoans. Lampreys are, together with hagfish, the sister group of jawed vertebrates (sharks to mammals), and the only living representatives of jawless vertebrates. Global mCG has been profiled in lamprey heart, brain, muscle and sperm, and was found to be highly heterogeneous at the majority of CpG sites, described as an “intermediate” between mosaic invertebrate mCG and vertebrate hypermethylation [32,33]. In addition, the lamprey genome contains a conserved mCG toolkit, including orthologues of DNMT1, DNMT3A, DNMT3B, DNMT3L, UHRF1, and TET proteins. Notably, lampreys as well as hagfish, only share one ancestral whole genome duplication (WGD) round with jawed vertebrates, which makes them unique in terms of this major genomic event when compared to both invertebrates and vertebrates [34–36]. Moreover, lampreys undergo a peculiar biological phenomenon termed programmed genome rearrangement (PGR), in which genomic DNA present in the germline is physically eliminated in somatic lineages during early embryogenesis, and is effectively entirely removed from the genome three days post fertilization [37– 41]. Homologues of genes in eliminated DNA sequences are enriched in functions related to germline development, and PGR is thought to prevent misexpression of genes with deleterious potential in somatic cells [42].

Here we sought to investigate whether the mCG configuration observed in the lamprey genome is compatible with embryonic epigenome remodeling processes, and whether 5mC may contribute to the targeting and elimination of germline-specific genes during PGR. To that end, we produced high resolution epigenome maps of sea lamprey development, employing whole-genome bisulfite sequencing (WGBS), biochemical identification of non-methylated DNA (BioCAP) [43] and Nanopore sequencing [44] of germline, embryonic, and adult somatic tissues. Our results demonstrate that lampreys undergo large scale maternal-to-paternal mCG remodeling. Unlike teleosts, however, where 5mC dynamics are localized to a relatively small number of defined loci, we discover developmental 5mC reprogramming occurring over partially methylated domains (PMDs) covering ∼30% of the entire genome, as well as at discrete gene regulatory elements. Furthermore, we show that prior to the onset of PGR, regions eliminated during early embryogenesis are pre-targeted by DNA methylation in sperm. Our results shed new light on mCG dynamics in early vertebrate lineages and provide important insights into the origins and evolution of vertebrate mCG reprogramming.

## RESULTS

### Disordered DNA methylation levels in embryonic and differentiated tissues

To investigate the extent of mCG reprogramming in the lamprey genome, we first questioned whether there are any observable mCG changes between diverse embryonic and adult lamprey tissues. We performed WGBS on seven lamprey samples comprising germline (sperm and egg), early embryonic (day 1 and day 2) and adult somatic tissues (brain, muscle, and peripheral blood mononuclear cells -PBMCs) in biological replicates (**Supplementary Figure S1, Supplementary Table S1**). The embryonic stages correspond to 64 cell (day 1) and pre-ZGA blastula (day 2), with ZGA occurring approximately 2.5-3 days post fertilization [41]. Initial analysis of the generated data revealed that all lamprey tissues display genomic mCG levels at an average of approximately 29-40% at the majority of CpG dyads, in clear contrast to the methylomes of both divergent and closely related vertebrate and invertebrate species, in which most CpG sites display high or low mCG (**Figure 1A-C**) [3,30]. These results are suggestive of considerable variation in mCG at the cellular level and are in close agreement with previously described adult lamprey datasets [32,33]. As other vertebrates, lamprey exhibits depleted (<1%) mCG in the mitochondrial genome (**Figure 1B**) and reduced non-CpG (CpH) genomic 5mC (**Figure 1D**). The exception to these trends is notable mCA enrichment (2.3%) in the brain, which was previously described as a deeply conserved epigenetic feature of the vertebrate neural system [33,45]. Altogether, the lamprey genome displays an intermediate DNA methylome state, in line with its phylogenetic position.

**Figure 1.**
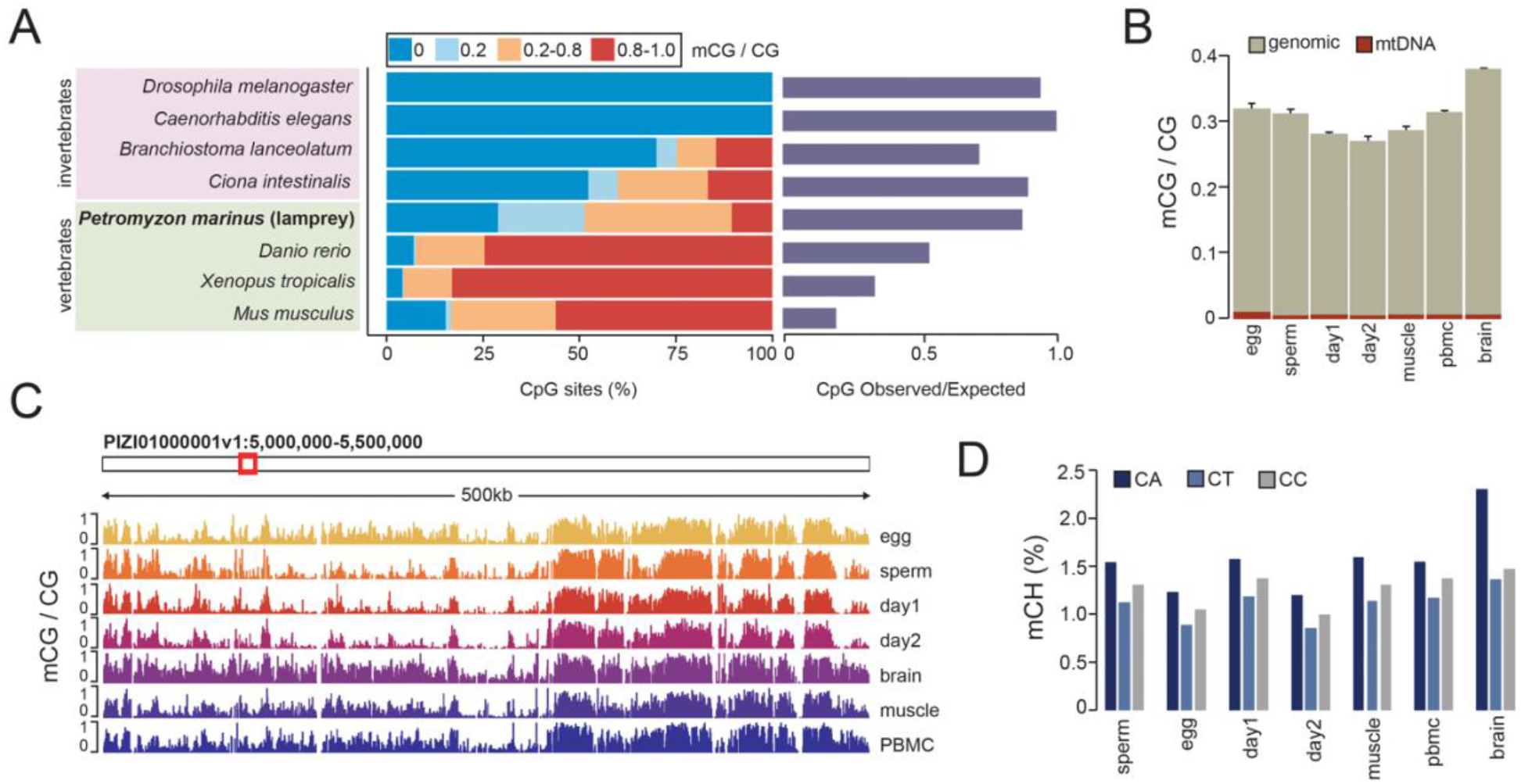
Global DNA methylation levels in seven lamprey tissues. **A)** Per CpG DNA methylation levels (mCG) and CpG observed/expected content for eight metazoan genomes. Global mCG levels obtained from publicly available WGBS datasets [6,98,99]. **B)** Genomic and mitochondrial mCG levels. **C)** Browser track depicting mCG profiles. **D)** Genomic mCH levels (%).

### Maternal-to-paternal embryonic reprogramming of partial DNA methylation

To better understand the origin and extent of intermediate DNA methylation in the sea lamprey genome, we partitioned the WGBS datasets into genomic blocks displaying highly disordered mCG levels (partially methylated domains, PMDs), and genomic regions located outside PMDs (non-PMDs) (**Figure 2A, Supplementary Tables S2-8**) [4,46]. To further interpret similarities between PMDs in each sample, we compared the genomic locations of PMDs between tissues and found an increased association between PMDs in; i) sperm, day 1 and day 2 methylomes, and in; ii) egg, brain, muscle and PBMC methylomes (**Figure 2B**). Overall, we found that ∼65% of the genome is classified as PMD in sperm, day 1 and day 2 tissues, which is significantly higher than what was observed in egg, brain, muscle and PBMC, where PMDs covered ∼46% of the genome (Welch two sample t-test, *p*-value < 0.001) (**Supplementary Figure S2A**). Finally, we found that while PMDs have largely similar mCG levels across tissues, notable differences in mCG levels could be observed between i) sperm, day 1 and day 2 non-PMDs, and ii) egg, brain, muscle, and PBMC non-PMDs (**Supplementary Figure S2B**). We next quantified the percentage of the genome that transitions from PMD to non-PMD and vice versa post-fertilization and found that ∼30% of the entire genome displays an altered PMD state in egg/sperm, egg/day 1 and egg/day 2 pairwise comparisons (**Figure 2C-D**). To better understand these transitions in DNA methylation states we calculated the proportion of discordant reads (PDR) [47,48] for both PMD groups, and plotted those against PMD mCG values (**Figure 2E**). The first group, comprising PMDs present in egg and other differentiated somatic tissues (brain, muscle, PBMCs, ∼4% of the genome), exhibited higher read discordancy and higher mCG levels in those tissues, when compared to sperm and embryonic tissues (day 1), where read discordancy (PDR) and mCG levels were notably lower (**Figure 2E, Supplementary Figure S2C**). The PMD group consisting of sperm, day 1, and day 2 samples (∼25% of the genome) displayed a notable difference in mCG levels when compared to egg, brain, muscle and PBMCs samples, even though the read discordancy was similar for this PMD group across all samples (**Figure 2E, Supplementary Figure S2C**). For both groups, changes in PMD state were paralleled by changes in mCG content, which we quantified at single CG sites (**Figure 2F**). For the first group, the developmental loss of PMDs was accompanied by a near complete loss of mCG (**Figure 2F, Supplementary Figure S2D**), whereas the second group, characterized by developmental PMD gain, displayed an increase in intermediate mCG levels (0.3 – 0.8 mCG/CG), with no apparent changes in low or high mCG fractions (**Figure 2F, Supplementary Figure S2D**). Finally, we also observed minor PMD transitions between sperm/day 1 and sperm/day 2; however, these represented a relatively lower proportion of the entire genome and were generally shorter than PMDs differentially identified between egg and sperm (**Figure 2C, Supplementary Figure S2E**). In summary, lamprey tissues can be separated into two major groups (sperm/day 1/day 2 and egg/brain/muscle/PBMC) based on their global DNA methylome structure, which is at least partly due to transitions in PMD states that may affect up to ∼30% of the genome.

**Figure 2.**
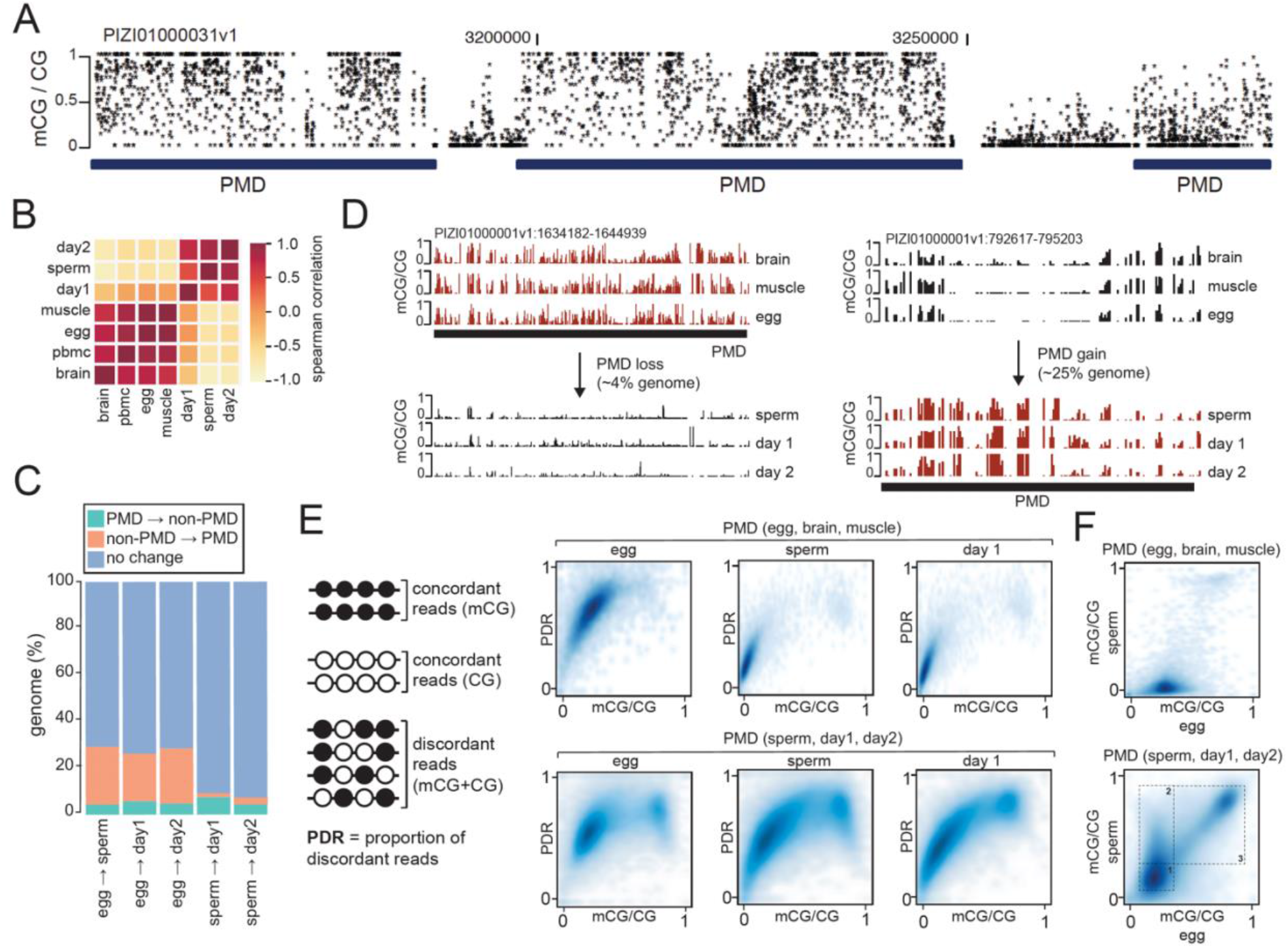
Reprogramming of partially methylated domains in the lamprey genome. **A)** Browser track depicting per CpG mCG levels for PMDs (dark blue) on scaffold PIZI01000031v1. **B)** Cluster heatmap of Spearman correlation of PMD overlap. **C)** Percentage of genome containing PMDs overlapping non-PMDs in pairwise comparisons of tissues. **D)** Browser track depicting mCG profiles of reprogrammed PMDs. **E)** Proportion of discordant reads (PDR) and mCG levels at reprogrammed PMDs. **F)** Pairwise comparisons of mCG levels at reprogrammed PMDs (1 – low mCG, 2 – intermediate mCG, 3 – high mCG).

### Differential DNA methylation enrichment at proximal and distal regulatory regions

Following observations that the lamprey epigenome could be partitioned into two major groups according to global tissue mCG patterning, we next wanted to explore whether similar distinctions could be observed at discrete regulatory regions. We focused this analysis on CpG islands (CGIs), also called non-methylated islands (NMIs), which are hypomethylated DNA sequences coinciding with vertebrate gene regulatory elements [49–52]. We experimentally identified non-methylated islands (NMIs) in six lamprey tissues (**Supplementary Table S9**) (sperm, day 1, day 2, brain, muscle, PBMCs) using BioCAP, a biochemical method based on protein affinity pulldown of unmethylated CpG-rich DNA [43]. As with the DNA methylome, we found that the NMI signal clustered into two major groups when compared between tissues: i) sperm, day 1, day 2, and ii) brain, muscle, PBMC (**Figure 3A**). This association was confirmed by PCA, (**Supplementary Figure S3A**) and by *k*-means clustering of mCG levels at NMIs merged from all examined tissues (**Figure 3B, Supplementary Figure S3B**). Moreover, we identified a core set of NMIs present in each tissue (*n* = 62,332) (**Supplementary Table S10**), and found DNA hypomethylation, increased CpG density and GC content, and association with accessible chromatin profiled in dorsal neural tube and whole heads [53], thus confirming that the chromatin and sequence features of lamprey NMIs resemble canonical vertebrate CGIs across multiple tissue types (**Figure 3C, Supplementary Figure S3C, D**). We next performed motif calling on core NMIs and found enrichment for transcription factor binding sites associated with ubiquitously active CGI-like promoters in both vertebrate [54,55] and invertebrate [56] genomes, including the methyl-sensitive transcription factor E2F, the enhancer box (E-box) regulatory motif and the transcription factor Nuclear Respiratory Factor (NRF) (**Figure 3D**). Furthermore, we found that NMIs were associated with ∼20-25% of all transcription start sites (TSS), underscoring their potential for gene regulatory function [50] (**Figure 3E, Supplementary Figure S3E**). Given the separation between NMIs in sperm, day 1, day 2 and brain, muscle, PBMC groups, we identified NMIs differentially enriched between these two groups based on normalized read density (false discovery rate < 0.05, fold-change > 2) to determine if grouped NMIs were found associated with specific genic features (**Supplementary Figure S3F**). Overall, we found a greater enrichment of brain/muscle/PBMC NMIs (*n* = 24,381) in genic regions compared to core NMIs and sperm/day 1/day 2 NMIs (*n* = 9,895), suggestive of differential mCG usage at NMIs in developmental and tissue-specific gene regulatory processes (**Figure 3F**).

**Figure 3.**
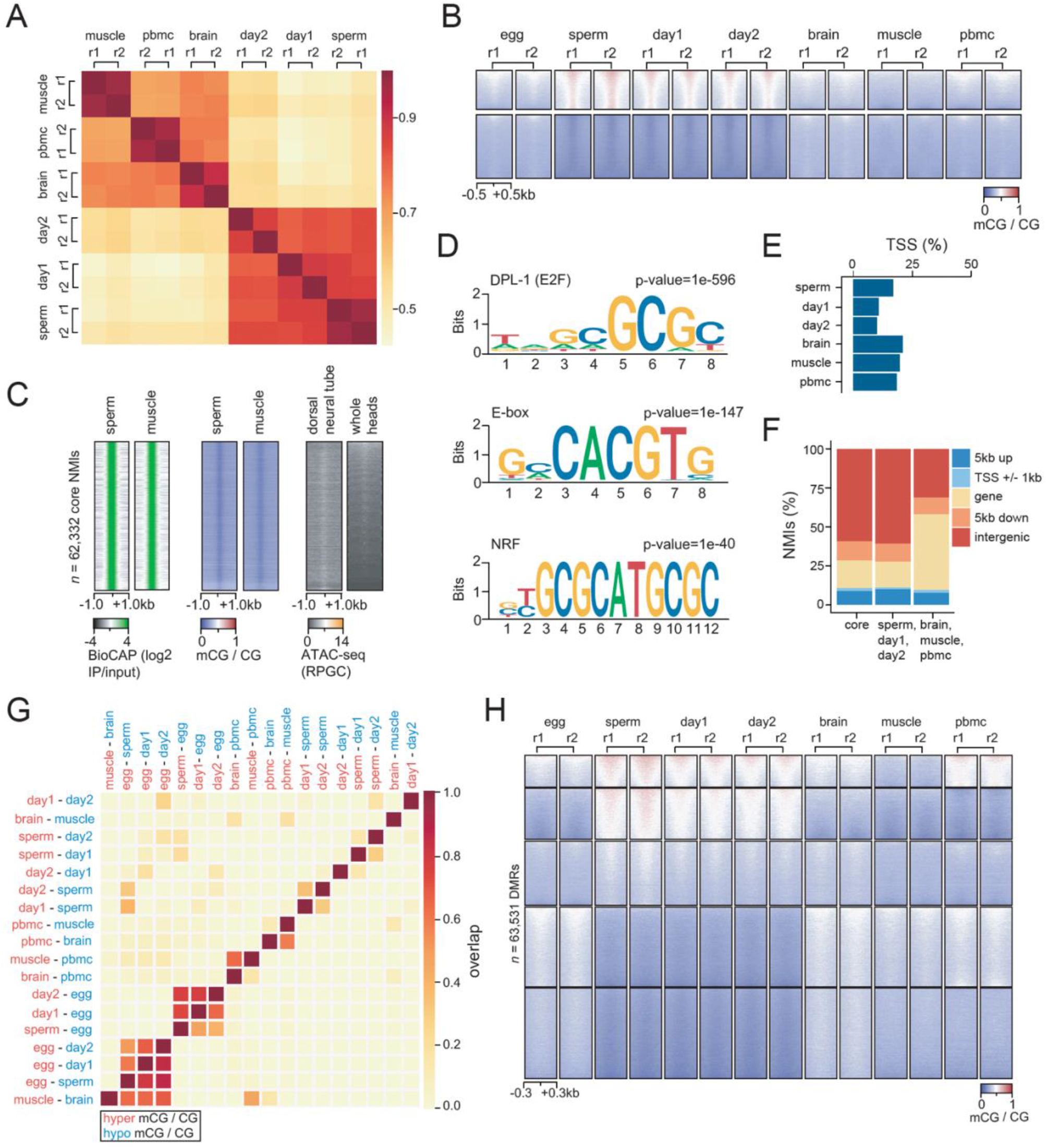
Developmental DNA methylation dynamics at *cis*-regulatory regions. **A)** Correlation heatmap of read density at NMIs. **B)** Heatmap of *k*-means clustered mCG signal at merged NMIs. **C)** BioCAP, mCG and ATAC-seq signal [53] at core NMIs. **D)** Sequence motif enrichment at core NMIs. **E)** Percentage of transcription start sites at protein-coding genes (*n* = 20,895) overlapped by NMI. **F)** Percentage of differentially enriched NMIs at genomic features. **G)** Cluster heatmap of DMR overlap. **H)** Heatmap of *k*-means clustered mCG signal at merged DMRs.

To obtain a more comprehensive view of developmental mCG dynamics in the lamprey, we performed pairwise comparisons of tissue-and stage-specific 5mC by identifying differentially methylated regions (DMRs) (ΔmCG > 0.2, p-value < 0.05). This approach resulted in the identification of ∼112,000 genomic regions displaying localized changes in methylation state (**Supplementary Table S11**). To determine tissue-specificity of DMRs, we assessed genomic co-localisation of all pairwise DMRs (**Figure 3G**). Overall, we found the greatest number of overlaps between egg/sperm, egg/day 1 and egg/day 2 DMRs, suggestive of maternal-to-paternal DMR reprogramming, as described in zebrafish [10,25– 28]. Next, we merged all DMRs into a single dataset and performed clustering of mCG levels (**Figure 3H**). mCG levels at DMRs grouped into two major tissue categories: i) sperm, day 1 and day 2; and ii) egg, brain, muscle and PBMC (**Supplementary Figure 4A-F**), in line with previous analyses. Altogether, we identified that DMRs (**Supplementary Table S11**) and differentially enriched NMIs represent ∼4% of the entire genome. While mCG patterns at these sequences recapitulate PMD dynamics, we found that ∼2% of the genome contains DMRs and differentially enriched NMIs located outside reprogrammed PMDs (**Supplementary Figure S4E**). This result indicates discrete 5mC remodeling occurring not only within, but also alongside large-scale genomic transitions in PMD state during early development.

### DNA sequences eliminated during PGR are hypermethylated in sperm

Programmed genome rearrangement (PGR) represents a unique biological mechanism for silencing deleterious gene loci [37–42]. During PGR in lamprey, DNA sequences containing several hundreds of genes are physically eliminated from the genome during early embryogenesis, effectively producing a somatic genome that is a smaller, reproducible fraction of the germline genome. PGR eliminates potential for somatic misexpression of germline-specific genes and is thus analogous to epigenetic silencing mechanisms described in vertebrate species, such as somatic gene silencing of cancer testis antigens [10,57–59]. As PGR is a crucial component of proper embryonic development in lamprey, we next interrogated the epigenetic landscape of sequences eliminated during PGR. We assembled whole-genome sequencing reads from blood and sperm [42] and called homozygous deletions in blood using CNVkit [60], identifying a total of 1,050 DNA sequences that we classified as eliminated sequences. We found that approximately 81% of regions identified by our analysis overlapped previously published eliminated sequences [42] (**Figure 4A-B, Supplementary Figure S5A**). Also, in line with previous reports, we observed that a considerable percentage (∼50%) of the eliminated fraction comprised diverse repetitive element classes, reflective of genomic DNA repeat content (**Supplementary Figure S5B**). We then selected 849 overlapping regions covering ∼9.3Mb of the genome as a stringent dataset for further genome-scale analyses. We used whole-genome sequencing [42] and WGBS read coverage to confirm that the presence of eliminated sequences was restricted to the germline (sperm) (**Figure 4C, Supplementary Figure S5C**). We also identified increased read density at eliminated sequences in day 1 and day 2 WGBS datasets, which is in line with previous findings that PGR occurs progressively during early development [37,40]. Finally, to confirm our findings using an orthogonal sequencing approach, we generated whole-genome Nanopore sequencing datasets for PBMCs and sperm in two biological replicates (**Supplementary Table S12**) and found increased read density at eliminated sequences in sperm, and read depletion in PBMCs (**Figure 4C, Supplementary Figure S5D**). Using our defined set of eliminated regions, we next set out to interrogate whether these genomic sequences exhibit a distinct mCG profile. Using sperm Nanopore sequencing data and sperm, day 1 and day 2 WGBS and BioCAP data, we identified DNA hypermethylation and depleted BioCAP signal at sequences eliminated during PGR (**Figure 4D, Supplementary Figure S5E**). We quantified mCG levels at these regions and found that the majority of CpG sites contained mCG > 75%, indicating a clear exception to global heterogeneous mCG (**Figure 4E** and **Supplementary Figure S5F**). Nevertheless, we could not identify any notable sequence features that would distinguish these targeted sequences from the genomic background (**Supplemental Figure S5G**), suggestive of sequence-independent mCG targeting. Altogether, our results suggest that mCG might play a role in the selection or perhaps even germline protection of eliminated DNA, thereby furthering our understanding of the molecular mechanisms governing PGR and underscoring the significance of mCG dynamics in diverse developmental processes.

**Figure 4.**
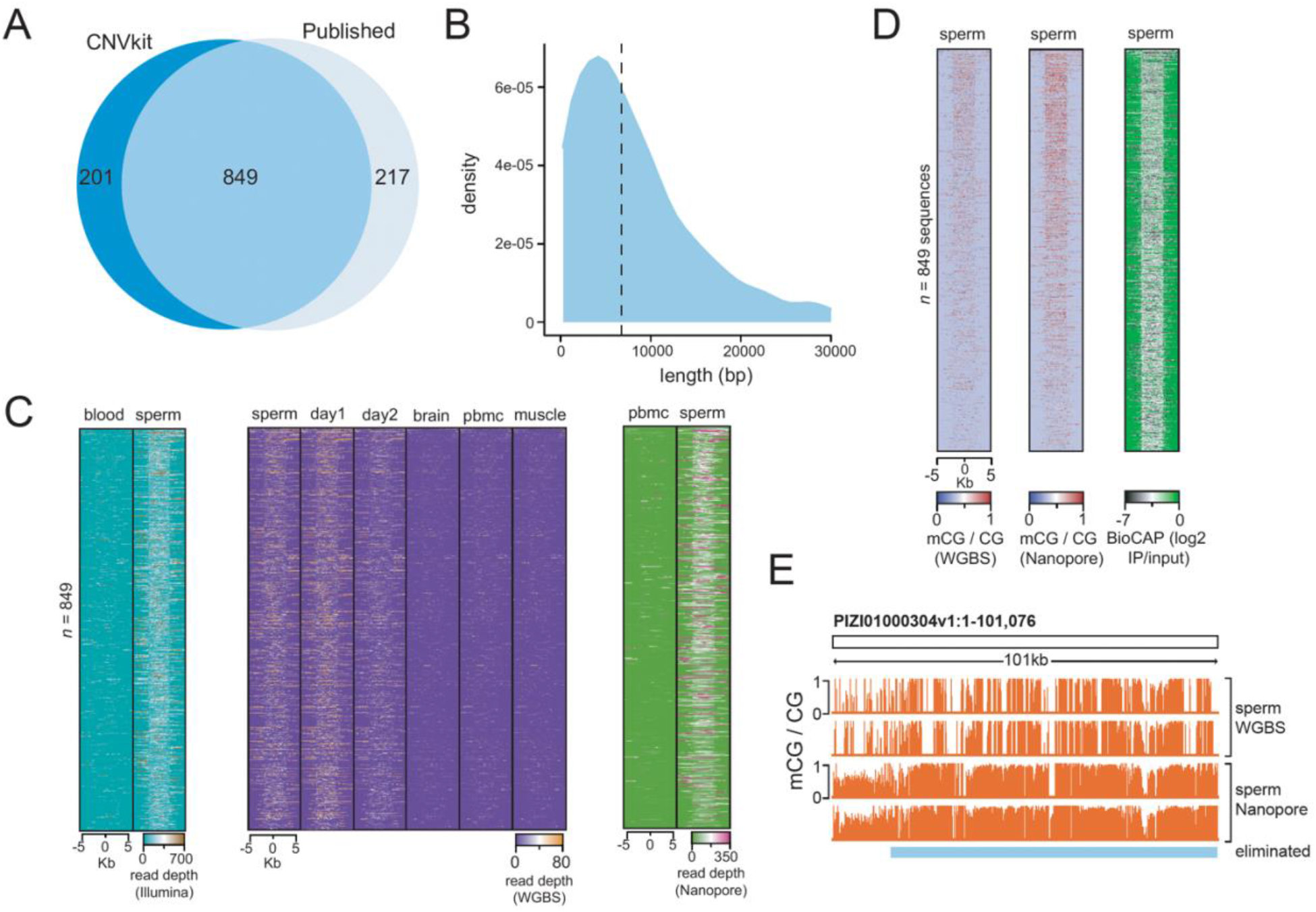
DNA methylation at eliminated sequences. **A)** Overlap of eliminated sequences previously published [42] and eliminated sequences identified using CNVkit. **B)** Distribution of lengths of eliminated sequences common to CNVkit and published data. **C)** Per-nucleotide read depth at eliminated sequences in whole-genome short-read [42], Nanopore, and bisulfite sequencing data. **D)** mCG and BioCAP signal at eliminated sequences. **E)** Browser track depicting mCG profiles from sperm (in biological replicate) at an eliminated sequence.

## DISCUSSION

The goal of this study was to identify and explore the dynamics and conservation of developmental 5mC reprogramming in the basal vertebrate sea lamprey. Lamprey represents an important model organism for studies of developmental 5mC function as it displays a distinctive, highly heterogeneous DNA methylome and undergoes genome-wide structural rearrangement during early embryogenesis [32,37–42]. Our study establishes for the first time that ∼30% of the Lamprey genome undergoes extensive developmental mCG reprogramming. We demonstrate that mCG patterns in the early embryo closely resemble sperm, thereby reflecting a maternal-to-paternal epigenome transition event upon fertilization. Moreover, we found that NMIs and hypomethylated DMRs in egg, brain, muscle and PBMCs overlap genic regions to a greater extent than those in sperm, day 1 and day 2, indicative of ZGA-associated epigenome remodeling [28]. These results therefore demonstrate that lamprey exhibits developmental mCG dynamics similar to zebrafish and medaka, thus providing valuable insights into the evolutionary origins of vertebrate mCG reprogramming [10,25–27] (**Figure 5**). Given this similarity in early developmental epigenome dynamics to teleost fish, it will be of interest to investigate to what extent mCG contributes to the development of body plans [61] and phenotypic plasticity [62,63] in the agnathan lineage.

**Figure 5.**
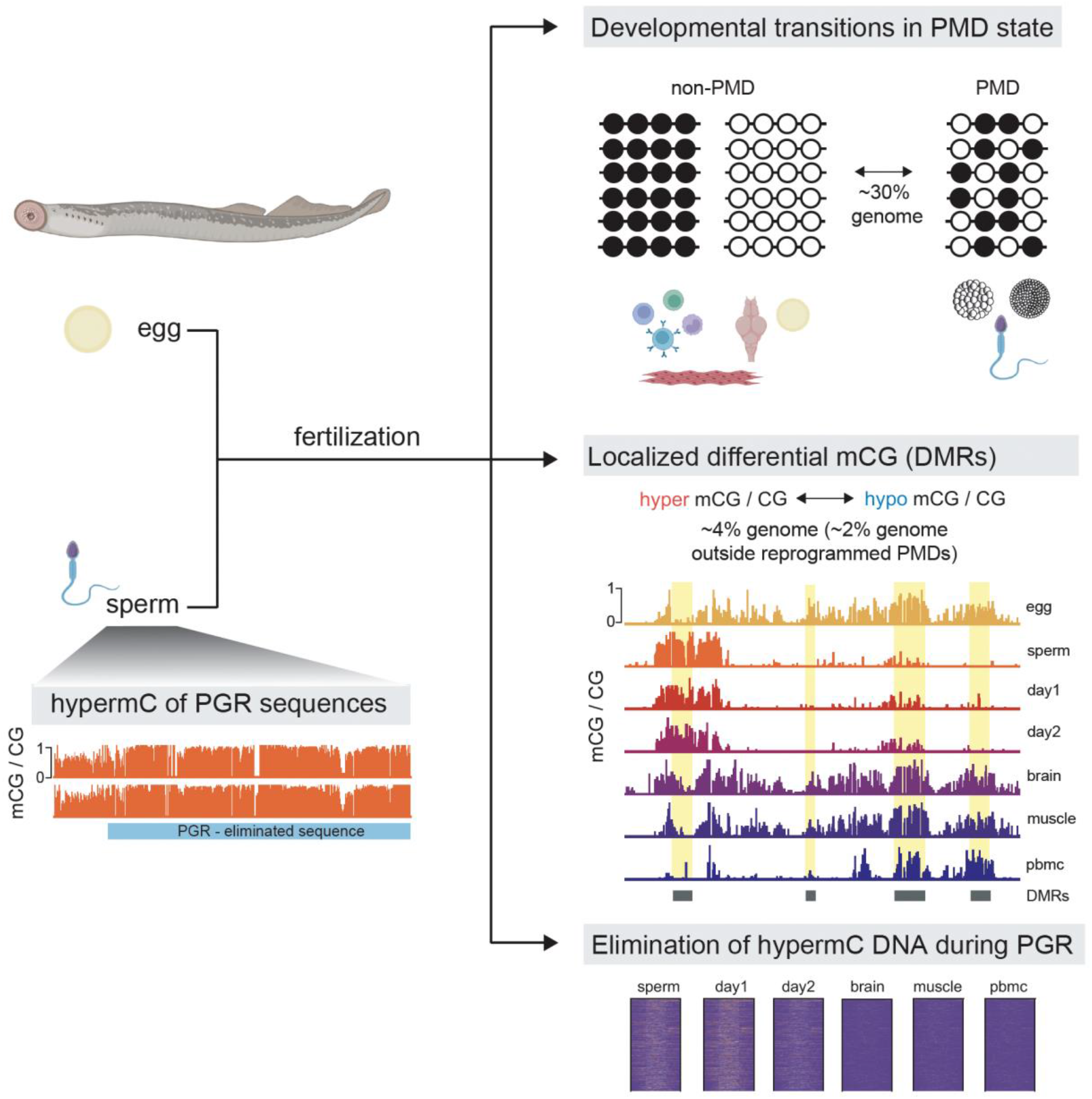
Overview of developmental mCG reprogramming in the sea lamprey. Early embryogenesis in the sea lamprey is characterized by maternal-to-paternal mCG reprogramming that affects up to 30% of the embryonic genome. These epigenetic remodeling events occur both on a large-scale (PMDs) and at discrete gene regulatory elements (DMRs, NMIs). Sequences that are eliminated during PGR in the early embryo are pre-targeted by DNA methylation in sperm.

Evolutionary studies of developmental 5mC dynamics will be useful in clarifying why vertebrate genomes shifted to hypermethylation as a default state during the invertebrate-to-vertebrate transition [64,65]. One plausible hypothesis is that genomic hypermethylation was developed as a mechanism to fine tune the increase in regulatory complexity arising from WGD events in vertebrate lineages [7–10,66]. It is currently thought that the sea lamprey genome underwent only a single duplication event, while gnathostome (jawed vertebrate) genomes are shaped by two rounds of WGD [34–36]. It is therefore possible that major epigenetic reprogramming observed during lamprey development is an ancient mechanism developed to balance genomic hypermethylation, which could be linked to vertebrate genome expansion. While our results suggest that maternal-to-paternal epigenome reprogramming is likely conserved in anamniotes, we observed notable differences between lamprey and previously profiled teleost species in the extent of these processes. In zebrafish, much of developmental 5mC reprogramming is limited to discrete gene regulatory elements, such as promoter CGIs [10,25–27]. In the lamprey however, while we identify changes to 5mC levels at putative regulatory elements, we also describe vast changes in genomic 5mC content between egg and sperm, with the sperm methylome being inherited in the early embryo. Approximately 30% of the lamprey genome transitions in PMD state post-fertilization (**Figure 5**), representing a large-scale developmental 5mC remodeling event in comparison to other anamniotes profiled to date. Moreover, we observed that maternal-to-paternal reprogramming of PMD states involves both developmental gain and loss of PMDs, even though developmental PMD gain before the ZGA onset was the predominant transition, affecting ∼25% of the genome.

It remains elusive exactly how mCG remodeling is achieved in lamprey. Based on mCG reprogramming dynamics identified in our study, it is not unlikely that lamprey utilize placeholder nucleosomes enriched in H2A.Z and H3K4me1 in a similar manner to zebrafish [28]. These placeholder nucleosomes are specific to the paternal germline in zebrafish and establish pre-ZGA chromatin in the early embryo. In zebrafish, placeholder nucleosomes are found at hypomethylated regulatory regions associated with developmental and housekeeping genes, where they deter DNMT activity while maintaining a transcriptionally quiescent state during cleavage stages [28]. Thus, it is plausible that NMIs in sperm and embryonic tissues represent sequences associated with a similar chromatin configuration in lamprey, which consequently display reduced mCG levels compared to tissues where such nucleosome positioning may not be present. In addition, H3K27me3 is a repressive chromatin modification deposited by Polycomb-group proteins, and is widely recognized to mediate silencing of developmental genes [67,68]. Some murine homologs of genes eliminated during PGR were found to be targets of Polycomb repressive complexes in mouse embryonic stem cells [42]. It will therefore be of particular importance to understand the genomic distribution and concentration of H3K27me3-marked regions in lamprey compared to vertebrate species that do not undergo PGR.

Our quantitative, base-resolution results of mCG targeting demonstrate that genomic regions eliminated during PGR are hypermethylated in sperm. This indicates that prior to fertilization and the onset of PGR, eliminated sequences are already epigenetically modified in the germline, which may facilitate accurate targeting and removal during PGR. Previous experiments performed using immunofluorescence assays suggested that mCG modification of targeted sequences could occur following their elimination and packaging into micronuclei [40]. It is important to note that the regions used in this study do not represent all sequences eliminated during PGR, due to technical challenges associated with the abundance of highly repetitive sequences in eliminated DNA [42]. PGR is not a lamprey-specific biological phenomenon; it has been described in several protozoan, invertebrate and vertebrate species, albeit with differences in timing and molecular mechanisms [69,70]. A common theme in PGR across diverse species is that eliminated sequences are associated with heterochromatin, such as in zebra finch [71], the ciliated protozoan *Tetrahymena* [72,73], and in sciarid flies [74,75]. Finally, an important question that remains open is how these regions remain refractory to elimination during germ cell development, which may be explained at least in part by the DNA hypermethylation observed in this study. Unlike mammals, fish do not undergo global mCG reprogramming in primordial germ cells, rather the paternal mCG configuration is retained [10,76]. If a similar mechanism would be observed in the lamprey, it would be plausible to suggest that DNA hypermethylation at eliminated sequences in sperm contributes to their retention in germline lineages. Further studies involving genomic localisation of DNMTs during lamprey embryonic development, including co-factors that are necessary for DNMT targeting to defined genomic locations, will deepen our understanding of the reprogramming processes described in this study.

In summary, we have demonstrated that despite high levels of mCG heterogeneity, the sea lamprey undergoes extensive maternal-to-paternal developmental DNA methylome remodeling predominantly associated with partially methylated DNA states. Moreover, we have shown that regions eliminated during PGR are marked by high mCG levels in sperm and pre-ZGA embryos. Altogether, this research demonstrates that the last common ancestor of vertebrates might have already presented extensive mCG reprogramming, and that this occurred before the second round of WGD that characterizes jawed vertebrates. Lampreys show lineage specific variations in the way in which they establish developmental mCG patterns, yet their epigenome configuration is suggestive of an early establishment of complex regulatory states that characterize the vertebrate lineage.

## MATERIALS AND METHODS

### Lamprey procedures

All lamprey embryonic and adult material was collected at the Hammond Bay Biological Station (Michigan, US). Embryos were obtained and grown as previously described [77]. All experimental procedures for culturing embryos were approved by Michigan State University Institutional Animal Care and Use Committee (AUF# 02/17-031-00). To produce sexually mature males and females for embryo fertilization, sea lamprey were transferred to the Ocqueoc River, Millersburg Michigan and held in cages (0.5m^3^) to allow natural sexual maturation in a riverine environment. Sea lamprey were checked daily for sexual maturity; sexually mature individuals were identified by applying abdominal pressure and checking milt expression or for ovulated oocyte expression. Sexually mature males and female lamprey were returned to HBBS and held until use for culturing lamprey embryos.

### Genomic DNA extraction from lamprey germline, embryonic and somatic tissues

Lamprey tissues were lysed in buffer containing 20 mM Tris, pH 8.0, 100 mM NaCl, 15 mM EDTA, 1% SDS and 0.5 mg/ml proteinase K for 3 h at 55 °C. Lysis was followed by two phenol:chloroform:isoamyl alcohol (25:24:1) extractions and subsequent centrifugation (5 min at 17,949g). DNA was then precipitated by adding 0.2 volumes of 4 M ammonium acetate and 3 volumes of 96% ethanol. The reaction was left on ice for a minimum of 30 min. The DNA precipitate was centrifuged for 20 min at 4°C (17,949g), and the pellet was washed with 500 μl of 70% ethanol and centrifuged for 5 min (17,949g) at room temperature. The pellet was then resuspended in 200 μl of TE buffer, and 1 μl of RNase A (20 μg/μl) was added. The reaction was left to proceed for 30 min at room temperature, after which time the DNA was precipitated with 0.1 volumes of 4 M ammonium acetate and 1 volume of isopropanol on ice for 2 h. The reaction was then centrifuged for 30 min at 4 °C (17,949g), and the pellet was resuspended in TE buffer.

### WGBS library preparation

WGBS was performed as described previously [10]. Briefly, DNA was spiked with unmethylated lambda phage fragments (Promega) and sonicated to approximately 300 base pairs. Sonicated DNA was concentrated in a vacuum centrifuge to a final volume of 20μL, required for bisulfite conversion with the EZ DNA Methylation-Gold Kit (Zymo Research). Bisulfite-converted DNA was then subjected to low input library preparation using the Accel-NGS Methyl-seq DNA Kit (Swift Biosciences). Briefly, the single-stranded, bisulfite-converted DNA was subjected to an adaptase reaction, followed by primer extension, adapter ligation (Methyl-seq Set A Indexing Kit-Swift Biosciences), and indexing PCR. Libraries were quantified using KAPA qPCR Library Quantification Kit (KAPA Biosystems) according to manufacturer instructions. Methylome libraries were sequenced on the Illumina HiSeqX platform (high-throughput mode, 150 bp, paired-end).

### WGBS read assembly

Read assembly was performed as described previously [10]. Briefly, sequenced reads in FASTQ format were trimmed using the fastp v0.12.5 tool (https://github.com/OpenGene/fastp) with the following settings: (fastp -i ${read_1} –I ${read_2} -o ${trimmed_read_1} –O ${trimmed_read_2} -f 10 -t 10 -F 10 -T 10). Trimmed reads were mapped (petMar3 genome reference, containing the lambda genome as chrL) using WALT with the following settings: -m 10 -t 24 -N 10000000 -L 2000 (https://github.com/smithlabcode/walt). PCR and optical duplicates were removed using Picard tools v2.3.0 (https://broadinstitute.github.io/picard/). Genotype and methylation bias correction was performed using MethylDackel software (https://github.com/dpryan79/MethylDackel). The number of methylated and unmethylated calls at each genomic CpG position were determined using MethylDackel (MethylDackel extract genome_lambda.fa $input_bam --mergeContext --minOppositeDepth 10 --maxVariantFrac 0.5 $input_bam -@ 32 --OT 0,0,0,0 --OB 10,0,0,0). Bisulfite conversion efficiency was estimated from the lambda phage spike-ins. Bedgraphs were converted to bigWig format using the bedGraphToBigWig script from kentUtils (https://github.com/ENCODE-DCC/kentUtils).

### WGBS methylome PCA

PCA of mCG levels was calculated using MethylKit PCASamples on two replicates of tissue methylomes using default parameters [78].

### Identification of partially methylated domains

Methylome replicates were merged and PMDs were called using MethylSeekR segmentPMDs function on scaffold PIZI01000001v1 using default parameters [46]. Tissue PMDs were intersected using Intervene pairwise (--compute frac) and Spearman correlation was found using the Intervene web application (https://asntech.shinyapps.io/intervene/) [79]. mCG levels from WGBS methylomes at PMDs and non-PMDs were found using bedtools intersect function [80]. To identify the percentage of discordant reads (PDR) at egg non-PMDs/sperm PMDs, per-read mCG status was determined from WGBS datasets using the MethylDackel perRead function. Discordant reads with at least four CpG sites were intersected with egg non-PMDs/sperm PMDs using bedtools coverage function (-counts) and mCG status was found by intersecting WGBS datasets with egg non-PMDs/sperm PMDs using bedtools map function (-o mean -null na). Scatterplots of PDR and mCG were generated using the smoothScatter function in R (nrpoints = 0). PMD genome coverage is calculated relative to the size of the petMar3 reference genome.

### BioCAP library preparation

BioCAP was performed on genomic DNA extracted from sperm, day 1, day 2, brain, muscle and PBMC in replicate and with input controls using published methods [81]. Briefly, genomic DNA was sonicated to approximately 200 base pairs and incubated with biotinylated zinc-finger CxxC protein domain from human KDM2B. NMIs were eluted in buffer containing 700mM-1M NaCl, 0.1% Triton X-100, 20 mM HEPES pH 7.9 and 12.5% v/v glycerol, then DNA was purified using the Wizard SV Gel and PCR Clean-Up System (Promega) according to manufacturer instructions. Library preparation was performed on BioCAP samples using TruSeq ChIP Sample Preparation (Illumina) according to manufacturer instructions. Libraries were amplified using 18 PCR cycles and quantified using KAPA qPCR Library Quantification Kit (KAPA Biosystems) according to manufacturer instructions. Libraries were sequenced on the Illumina NovaSeq 6000 (150 bp, paired-end), generating 33M-307M read pairs per sample.

### BioCAP read assembly

BioCAP reads were trimmed using trimmomatic (ILLUMINACLIP:TruSeq3-PE-2.fa:2:30:10 SLIDINGWINDOW:5:20 LEADING:3 TRAILING:3 MINLEN:25) [82] and aligned to the petMar3 reference genome using bowtie2 (-p 10 - N 1 --very-sensitive -X 2000 --no-mixed --no-discordant) [83]. The resulting alignments in BAM format were deduplicated using Picard MarkDuplicates (REMOVE_DUPLICATES=true). NMIs were identified using the macs2 callpeak function [84]. When replicate is not specified, dataset refers to NMIs common to both replicates. Bigwig (log2 IP/input) files were generated using deepTools bamCompare (--ignoreDuplicates --binSize 50 -- centerReads --extendReads) [85].

### ATAC-seq read assembly

Publicly available ATAC-seq data in fastq format was downloaded [53]. ATAC-seq reads were trimmed using trimmomatic (ILLUMINACLIP:TruSeq3-PE-2.fa:2:30:10 SLIDINGWINDOW:5:20 LEADING:3 TRAILING:3 MINLEN:25) and aligned to the petMar3 reference genome using bowtie2 (-p 10 - N 1 --very-sensitive -X 2000 --no-mixed --no-discordant). To include only nucleosome-free read alignments, alignments exceeding 100bp were removed. Aligned reads were deduplicated using Picard MarkDuplicates (REMOVE_DUPLICATES=true). Bigwig (RPGC) files were generated using deepTools bamCoverage (--normalizeUsing RPGC -- numberOfProcessors 4 --ignoreDuplicates -- binSize 50 --centerReads --extendReads -- effectiveGenomeSize 1040808129).

### Epigenetic features of NMIs

Clustering and PCA of NMI read counts was performed using DiffBind dba.count and dba.plotPCA functions, respectively [86]. To calculate mCG clustering, all NMI datasets were concatenated and overlapping sequences were merged using bedtools merge function. Heatmaps of mCG signal at merged NMIs were generated using deepTools computeMatrix function (computeMatrix reference-point --referencePoint center --binSize 10 --afterRegionStartLength 500 - -beforeRegionStartLength 500), with replacement of NaN values with mean mCG (0.33) after the matrix file was generated, followed by the plotHeatmap function (--kmeans 5 --yMin 0 -- yMax 1). NMIs present in every tissue were found by intersecting all NMI datasets using bedops intersect function [87]. Heatmaps of BioCAP signal at core NMIs were generated using deepTools computeMatrix function (computeMatrix reference-point --referencePoint center --binSize 10 --afterRegionStartLength 1000 --beforeRegionStartLength 1000 -- missingDataAsZero), followed by the plotHeatmap function (--sortRegions no). Heatmaps of mCG signal at core NMIs were generated using deepTools computeMatrix function (computeMatrix reference-point --referencePoint center --binSize 10 --afterRegionStartLength 1000 --beforeRegionStartLength 1000), with replacement of NaN values with mean mCG for each tissue after the matrix file was generated, followed by the plotHeatmap function (-- sortRegions no). Heatmaps of ATAC-seq signal at core NMIs were generated using deepTools computeMatrix function (computeMatrix reference-point --referencePoint center –binSize 10 --afterRegionStartLength 1000 -- beforeRegionStartLength 1000 -- missingDataAsZero), followed by the plotHeatmap function (--sortRegions no).

### Sequence motifs at core NMIs

Enriched sequence motifs at core NMIs were found using Homer findMotifsGenome.pl function (-size 200) with default parameters [88]. Sequence logos were visualized using the ggseqlogo package in R [89]. CpG observed/expected and GC content was calculated using bedtools getfasta and an in-house generated R script.

### Localization of NMIs at genomic features

NMIs differentially enriched in sperm/day 1/day 2 and brain/muscle/PBMC were found using DiffBind dba.analyze function (false discovery rate < 0.05, fold change > 2). petMar3 gene predictions were downloaded [42] and overlap with NMIs was identified using bedtools intersect function. Barplots were generated using ggplot2 geom_bar function in R [90]. Heatmaps of BioCAP signal at TSS were computeMatrix generated function using deepTools (computeMatrix reference-point --referencePoint center –binSize 10 --afterRegionStartLength 1000 -- beforeRegionStartLength 1000 -- missingDataAsZero), followed by the plotHeatmap function (--sortRegions no).

### Differentially methylated regions

DMRs were called from WGBS data with the DSS software [91,92] with the following parameters: delta=0.2, p.threshold=0.05, minlen=100, minCG=10, dis.merge=100. DMRs were intersected using Intervene pairwise (--compute frac) and the heatmap was generated using the Intervene web application (https://asntech.shinyapps.io/intervene/). To calculate mCG clustering, all DMRs were concatenated and overlapping sequences were merged using bedtools merge function. Heatmaps of mCG signal at merged DMRs were generated using deepTools computeMatrix function (computeMatrix reference-point --referencePoint center --binSize 10 --afterRegionStartLength 300 - -beforeRegionStartLength 300), with replacement of NaN values with mean mCG (0.33) after the matrix file was generated, followed by the plotHeatmap function (--kmeans 5 --yMin 0 -- yMax 1). mCG levels at DMRs were found using bedtools intersect function and histograms of mCG at DMRs were created using ggplot2 geom_histogram function. Heatmaps of BioCAP signal at DMRs were generated using deepTools computeMatrix function (computeMatrix reference-point --referencePoint center –binSize 10 --afterRegionStartLength 300 -- beforeRegionStartLength 300 -- missingDataAsZero), followed by the plotHeatmap function (--sortRegions no). CpG density was calculated using bedtools getfasta and an in-house generated R script.

### Localization of DMRs at genomic features

petMar3 gene predictions were downloaded [42] and overlap with DMRs was identified using bedtools intersect function. Barplots were generated using ggplot2 geom_bar function in R.

### Whole-genome sequencing read assembly

Publicly available blood and sperm whole-genome sequencing reads in fastq format were downloaded [42]. Reads were trimmed using trimmomatic (ILLUMINACLIP:TruSeq3-PE-2.fa:2:30:10 SLIDINGWINDOW:5:20 LEADING:3 TRAILING:3 MINLEN:25) and aligned to the petMar3 reference genome using bowtie2 (-p 10 - N 1 --very-sensitive -X 2000 --no-mixed --no-discordant). Reads mapping to multiple genomic regions were removed using samtools [93].

### Nanopore library preparation and sequencing

Library preparation was performed using ONT ligation sequencing (SQK-LSK110). Samples were sequenced across two PromethION (FLO- PRO002) flow cells for 72 hours.

### Nanopore read assembly and mCG calling

Raw ONT sequencing data was converted to BLOW5 format [94] using slow5tools v0.3.0 [95] then base-called using Guppy v5.0.13 (high-accuracy model). Resulting fastq files were aligned to the petMar3 reference genome using minimap2 v2.22 [96]. 5mC profiling on CpG sites within petMar3 was performed using f5c v0.7 [97] with BLOW5 data input. CpG methylation frequencies were determined using the meth-freq tool in f5c.

### Identification of sequences eliminated during PGR

CNVkit (-m wgs) was used to identify sequences eliminated during PGR [60]. We defined homozygous deletions as sequences with a copy number of 0 in blood compared to sperm. Sequences were validated using coverage metrics from short-read whole-genome sequencing, WGBS and Nanopore sequencing. Read depth was calculated using samtools depth function and converted to bigWig format using the bedGraphToBigWig script from kentUtils. Heatmaps of read coverage at eliminated sequences were generated using deepTools computeMatrix function (computeMatrix scale-regions -- regionBodyLength 7000 --missingDataAsZero -- binSize 10 --afterRegionStartLength 5000 -- beforeRegionStartLength 5000), followed by the plotHeatmap function (--sortRegions no).

### mCG at eliminated sequences

Heatmaps of mCG signal at eliminated sequences were generated using deepTools computeMatrix function (computeMatrix reference-point -- regionBodyLength 7000 --binSize 10 -- afterRegionStartLength 5000 -- beforeRegionStartLength 5000), with replacement of NaN values with mean mCG for each sequencing technique after the matrix file was generated, followed by the plotHeatmap function (- -sortRegions no). Heatmaps of BioCAP signal at eliminated sequences were generated using deepTools computeMatrix function (computeMatrix reference-point --binSize 10 -- regionBodyLength 7000 –afterRegionStartLength 5000 --beforeRegionStartLength 5000 -- missingDataAsZero), followed by the plotHeatmap function (--sortRegions no). mCG status at eliminated sequences was found by intersecting WGBS datasets with eliminated sequences using bedtools intersect function and violin plots were generated using ggplot2 geom_violin function in R. Repeat content was found by intersecting the petMar3 RepeatMasker track with eliminated sequences using bedtools intersect function. Dinucleotide frequency at eliminated sequences was calculated using the faCount script from kentUtils.

### Availability of data and materials

BioCAP-seq and WGBS data generated in this study are available from NCBI Gene Expression Omnibus (GEO SuperSeries accession number GSE220553). Nanopore sequencing data generated in this study is available from NCBI (BioProject accession number PRJNA783432). ATAC-seq data used in this study was downloaded from GSE112072.

## Supporting information

Angeloni_et_al_2023_Supplementary_Material

Angeloni_et_al_2023_Tables_S1-S12

## Authors’ contributions

OB conceived the study. Lamprey embryo and adult tissue collection and DNA extraction was performed by OB, XZ, WL and SF. Sequencing libraries were generated by AA. CxxC protein domain from human KDM2B used for BioCAP was prepared by DK and RJK. Nanopore library preparation and sequencing was performed by

JMH. Nanopore read assembly and 5mC calling was performed by HG and IWD. AA performed all other data analysis. OB participated in WGBS data analysis. AA and OB wrote the manuscript. All authors contributed to, read, and approved the final manuscript.

## Acknowledgements

Figures were created with the help of BioRender.com software. The authors thank Alex de Mendoza for critical reading of the manuscript.

