## Supplementary material for "Extensive DNA methylome rearrangement during early lamprey embryogenesis": Angeloni_et_al_2023_Supplementary_Material

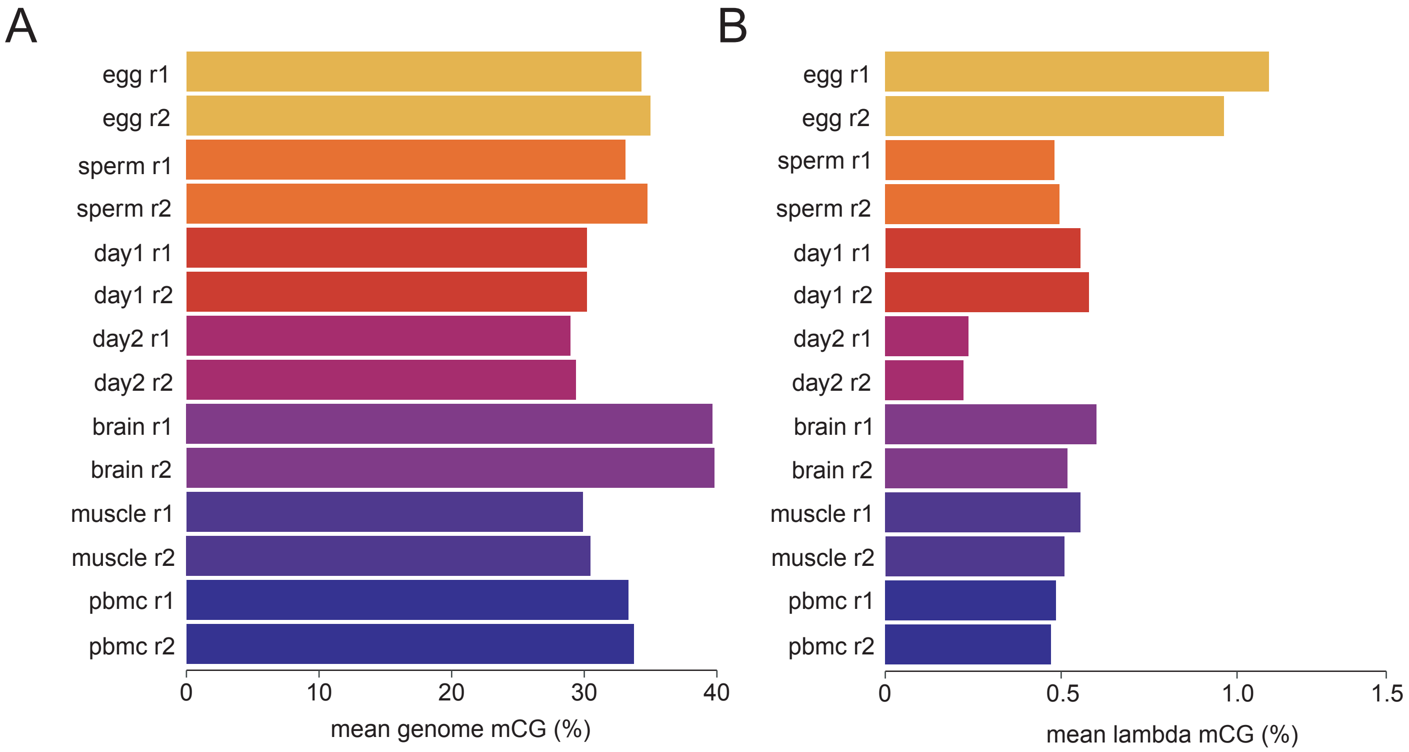


**Supplementary Figure S1.** Global DNA methylation levels in WGBS biological replicates. **A)** Mean genomic mCG levels. **B)** mCG levels in the unmethylated lambda phage DNA spike-in control for every WGBS dataset generated for this study.

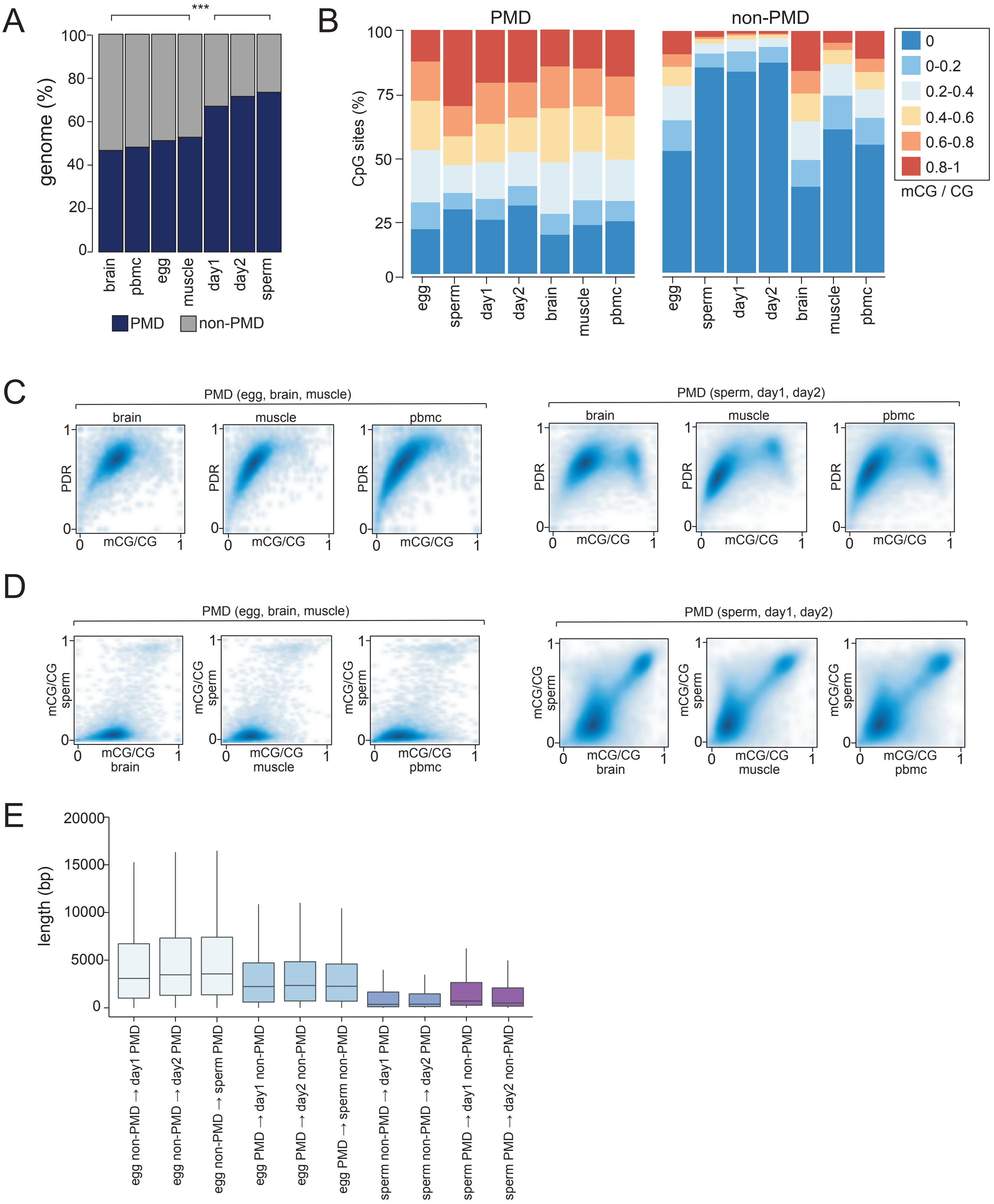


**Supplementary Figure S2.** Sequence features of lamprey PMDs**. A)** Percentage of the lamprey genome covered by PMDs (dark blue) and non-PMDs (grey) in embryonic and adult somatic and germline tissues. Welch two sample t-test, p-value < 0.001. **B)** Percentage of CpG sites displaying low (0, 0-0.2), intermediate (0.2 - 0.4, 0.4 - 0.6, 0.6 - 0.8) and high (0.8 - 1.0) mCG levels at PMDs and non-PMDs in embryonic and adult somatic and germline tissues. **C)** Proportion of discordant reads (PDR) and mCG levels for adult somatic tissues at reprogrammed PMDs. **D)** Pairwise comparisons of mCG levels at reprogrammed PMDs. **E)** Distribution of sequence lengths of developmentally reprogrammed PMDs.

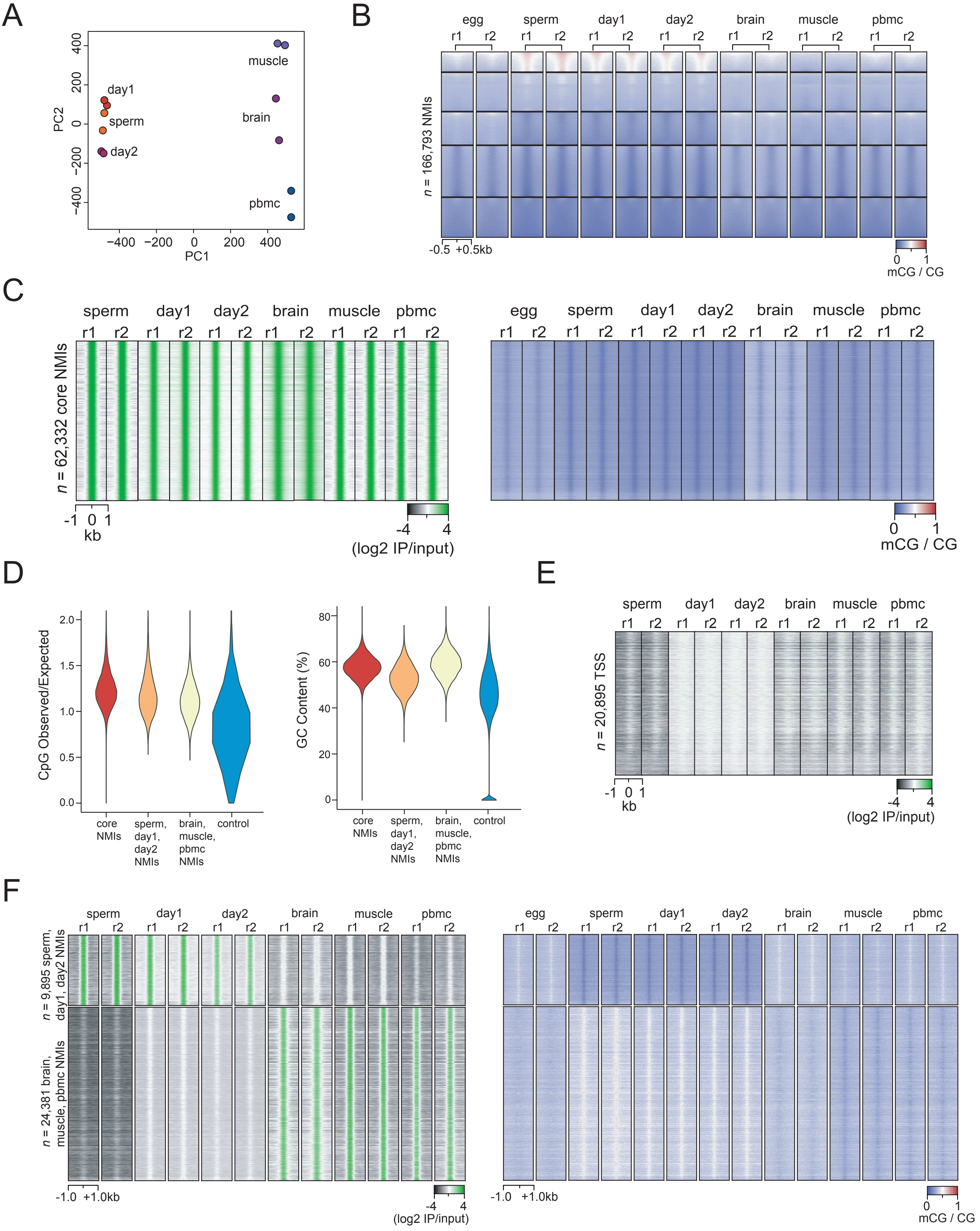


**Supplementary Figure S3. Sequence and epigenetic features of shared and tissue-specific NMIs. A)** PCA of normalized BioCAP read density at NMIs. **B)** *K*-means clustered mCG signal at merged NMIs. **C)** BioCAP and mCG signal at core NMIs. **D)** Distribution of CpG observed/expected ratio and GC content at: core NMIs; NMIs enriched in sperm, day 1 and day 2; NMIs enriched in brain, muscle and PBMC; and random control sequences. **E)** BioCAP signal at transcription start sites of protein-coding genes. **F)** BioCAP and mCG signal at NMIs enriched in sperm, day 1 and day 2, and at NMIs enriched in brain, muscle and PBMC.

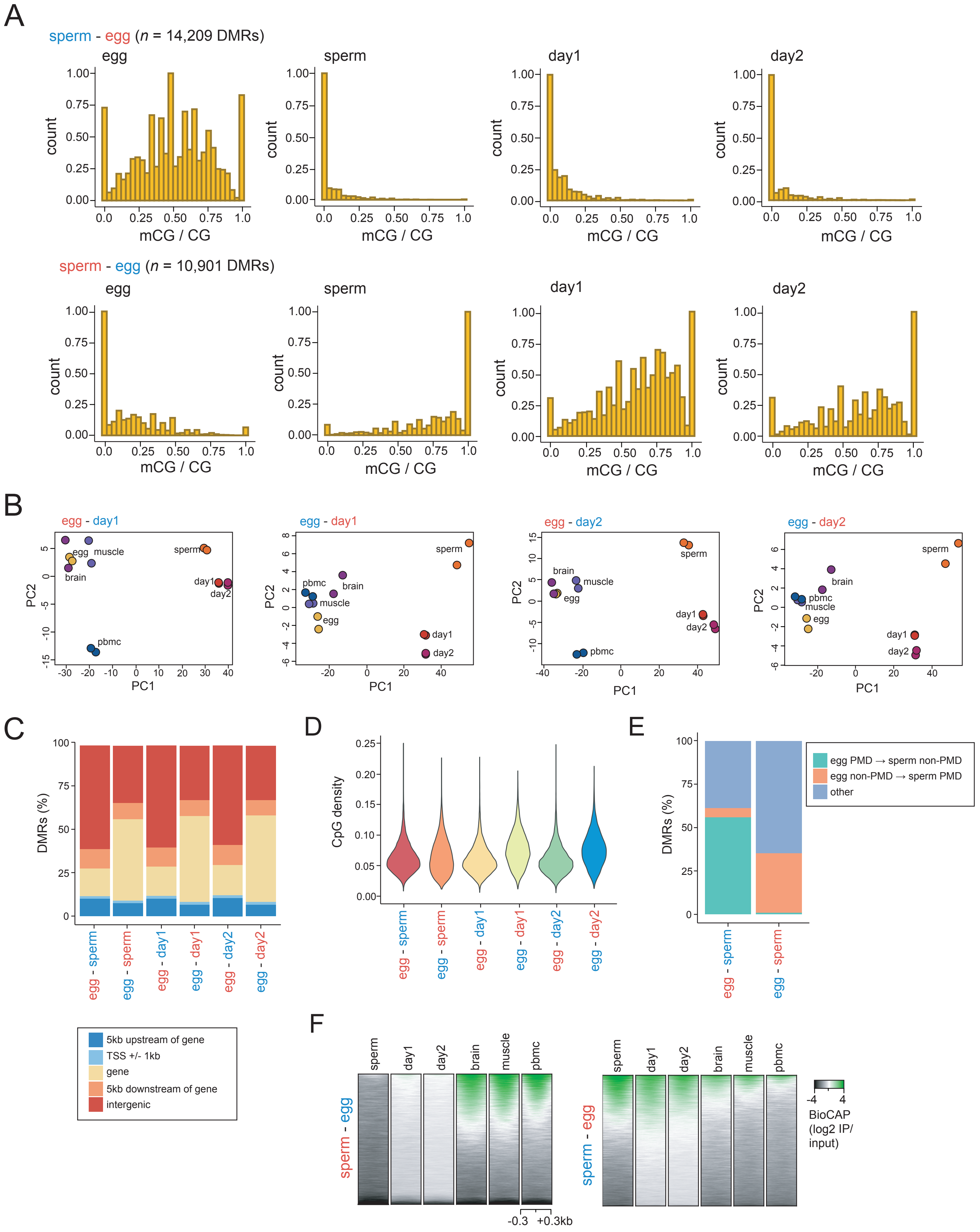


**Supplementary Figure S4. Maternal to paternal reprogramming at differentially methylated regions (DMRs) during lamprey embryogenesis. A)** Distribution of mCG levels at egg/sperm DMRs. **B)** PCA of mCG levels from WGBS data (in biological replicate) at egg/day 1 and egg/day 2 DMRs. **C)** Percentage of egg/sperm, egg/day 1 and egg/day 2 DMRs overlapping diverse genomic features. **D)** Distribution of CpG density at egg/sperm, egg/day 1 and egg/day 2 DMRs. **E)** Percentage of egg/sperm DMRs overlapping egg/sperm PMDs. **F)** BioCAP signal at egg/sperm DMRs. BioCAP signal is depicted in descending order of signal intensity.

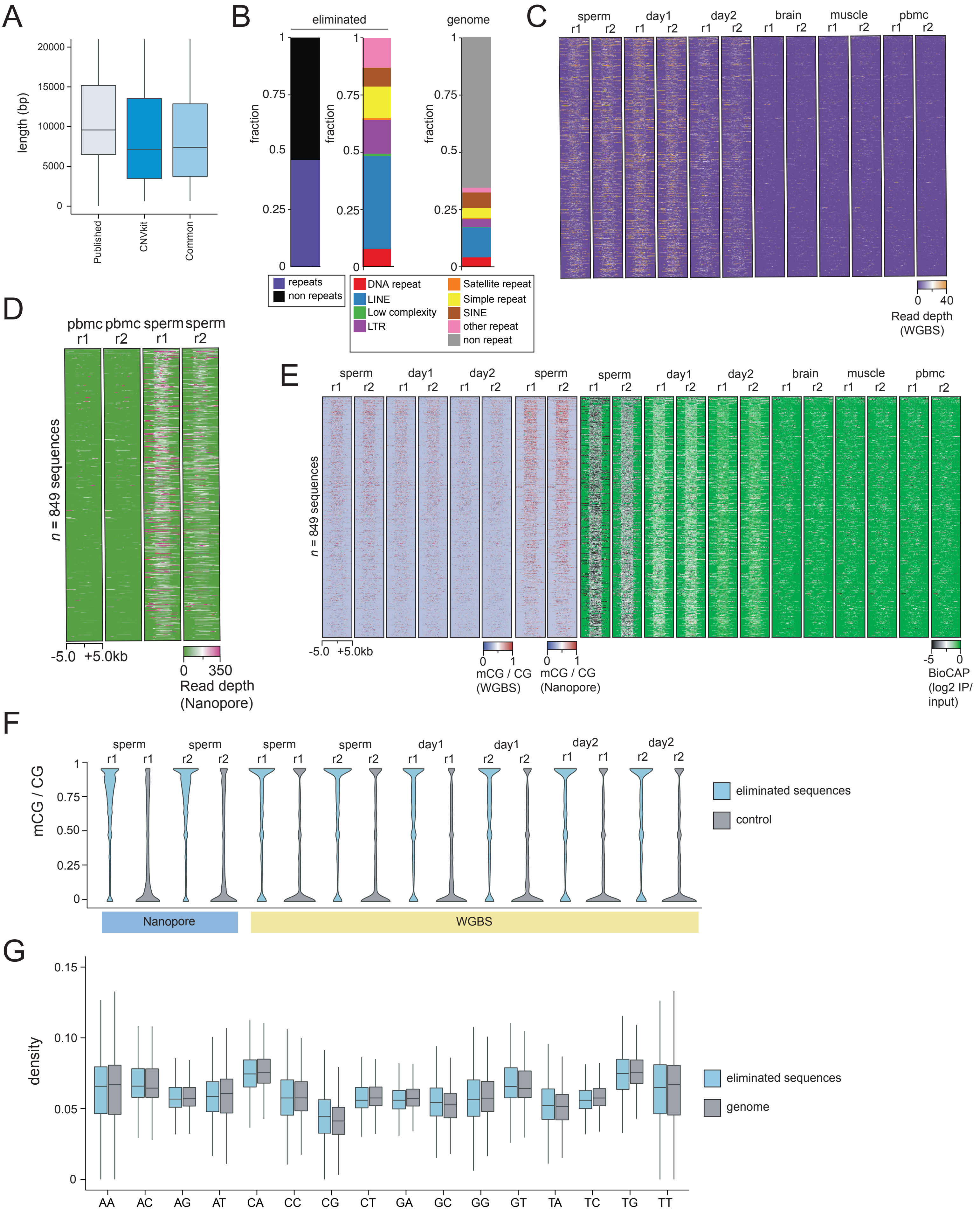


**Supplementary Figure S5. Sequence and epigenetic features of eliminated regions. A)** Distribution of lengths of previously published eliminated DNA sequences [42], eliminated sequences detected using CNVkit, and eliminated sequences common to both methods. **B)** Repeat content of eliminated DNA sequences (left), repeat composition as per Repeatmasker track of eliminated DNA sequences (middle), and genomic percentage and composition of repetitive DNA sequences (Repeatmasker) in the petMar3 reference genome. **C)** Per-nucleotide read depth at eliminated sequences in bisulfite sequencing data. **D)** Per-nucleotide read depth at eliminated sequences calculated from Nanopore sequencing data. **E)** mCG and BioCAP signal (in biological replicates) at eliminated DNA sequences. **F)** mCG levels (in biological replicate) at eliminated and control sequences from Nanopore and WGBS data. **G)** Dinucleotide frequencies at eliminated sequences and genome-wide.
